# Inoculum and disease dynamics of citrus greasy spot caused by *Zasmidium citri-griseum* in sweet orange in Panama

**DOI:** 10.1101/2022.08.26.505504

**Authors:** Vidal Aguilera-Cogley, Elena Sedano, Antonio Vicent

**Affiliations:** Grupo de Investigación de Protección Vegetal, Centro de Innovación Agropecuaria Divisa, Instituto de Innovación Agropecuaria de Panamá (IDIAP), 0619 Herrera, Panamá; Centre de Protecció Vegetal i Biotecnologia, Institut Valencià d’Investigacions Agràries (IVIA), 46113 Moncada, Spain

**Keywords:** autoregressive temporal component, Central America, epidemiology, INLA, *Mycosphaerella citri*

## Abstract

Citrus greasy spot, caused by *Zasmidium citri-griseum*, is a disease characterized by inducing premature defoliation and a reduction in yield in different citrus species. Greasy spot is the most prevalent fungal disease in sweet orange in Panama. Nevertheless, no epidemiological information is available. In this study, the dynamics of the defoliation, inoculum production, airborne inoculum, and infection periods of *Z. citri-griseum* and their associations with environmental conditions were determined in Panama. The period from December to April was characterized by greater defoliation of trees, with the greatest amount of leaf litter being produced in January and February. The number of days until total leaf decomposition (*DLD*) was related to the number of rainy days >1mm (*NRD*), accumulated rainfall (*AR*), and average relative humidity (*ARH*). The number of ascospores released from leaf litter (*ASCL*) was related to *DLD, NRD, AR*, and average temperature (*AT*). The greatest amounts of airborne ascospores (*AASC*) of *Z. citri-griseum* occur during April and May, when the rainy season begins in Panama. Similarly, the highest incidence (*INC*) of greasy spot in the trap plants coincided with the months of the greatest availability of airborne ascospores. However, infections were also recorded during other times of the year. The *AASC* or *INC* data were fitted to Bayesian models including meteorological variables and an autoregressive temporal component, the latter being the most influential. The results obtained in this study will allow the development of more efficient and sustainable fungicide programs for greasy spot control in Panama.

## 1 INTRODUCTION

Sweet orange (*Citrus sinensis* [L.] Osbeck) is an important fruit, widely cultivated in Central America. In this region, the production of sweet orange is estimated at 5,638,789 tonnes, with a total cultivated area of 393,826 ha (FAOSTAT 2022). In Central America, sweet orange is grown in tropical climate conditions, with large amounts of rainfall that favor the development of important fungal diseases associated with yield and quality losses, such as citrus greasy spot, caused by *Zasmidium citri-griseum* (F.E. Fisher) U. Braun and Crous (=*Mycosphaerella citri* Whiteside) (Aguilera-Cogley and Vicent 2019; Mondal and Timmer 2006).

Sweet orange has become the main species of citrus in Panama with a total cultivated area of 14,875 ha, which represents 98% of the total citrus-growing areas in the country (MIDA 2021). The crop develops under a tropical climate, with mean temperatures generally above 18°C and a relatively long rainy season from May to November, and a dry season from December to April (Peel et al. 2007; ETESA 2022). As in other tropical areas, sweet orange in Panama is typically rain-fed, with a major bloom in May to June followed by a minor bloom in September to October.

The environmental conditions in Panama are highly favorable for the development of greasy spot on sweet orange, which represents a limiting factor for production in most citrus-growing areas in the country. Recent surveys have indicated that greasy spot is the most prevalent fungal disease in sweet orange in Panama, with 67.2% of the orchards affected (Aguilera-Cogley and Vicent 2019). Greasy spot induces premature defoliation and a reduction in yield in different citrus species. Most epidemiological studies on this disease have been conducted in the USA, in citrus areas in Florida and Texas, where the disease has historically been a problem (Timmer et al. 1995; Timmer et al. 1980; Timmer et al. 2000; Whiteside 1970). In Central America, epidemiological studies on greasy spot have been carried out only in Costa Rica (Hidalgo et al. 1997).

Ascospores formed in pseudothecia on decaying leaves on the orchard floor are the primary inoculum source of *Z. citri-griseum* (Mondal and Timmer 2003; Timmer and Gottwald 2000). Ascospores are discharged in response to wetting and dispersed by wind currents (Timmer 1999). Once deposited on the leaf surface, ascospores germinate and the resulting mycelia grow epiphytically (Timmer et al. 2004). Conidia of *Z. citri-griseum* have also been reported, but they are considered of minor epidemiological relevance (Timmer and Gottwald 2000).

Trees affected by greasy spot typically show yellow mottle on the adaxial surface of leaves and yellow-brown, slightly raised pustules on the abaxial surface. The disease is more severe on grapefruit, lemons, and early-maturing sweet oranges than on late-maturing sweet oranges and mandarins (Timmer et al. 2003). Greasy spot is characterized by a relatively long incubation period and foliar symptoms are usually observed in mature leaves. Premature leaf drop may render the affected trees almost completely defoliated, thereby reducing yield and fruit size (Mondal and Timmer 2006; Timmer and Gottwald 2000). Excessive defoliation of citrus trees due to greasy spot occurs before the end of the winter, and then the development of the spring flush is impaired and fruit yield is reduced (Whiteside 1977).

The production of ascospores of *Z. citri-griseum* usually occurs several weeks after a major leaf drop, depending on rain events and temperature (Timmer et al. 2004). Rainfall or micro-sprinkler irrigations are also important for ascospore release (Mondal et al. 2003; Whiteside 1974). In Florida, the dynamics of ascospore release varied over the years. Initially, Whiteside (1974) reported that the highest concentration of ascospores in the air occurred in June. More recently, with the increase in the frequency of micro-sprinkler irrigation the peak of ascospore release has moved forward to April and May (Timmer and Gottwald 2000). In Texas, where rains begin later, the peak of ascospore release occurs in July-September (Timmer et al. 1980). Infections mainly depend on how fast the epiphytic mycelium develops on the leaf surface, which is influenced by temperature and rainfall (Timmer et al. 2004). In Costa Rica, where the climatic conditions are different from those in Florida and Texas, the peak of ascospore release occurred in June and declined rapidly in July (Hidalgo et al. 1997).

The infection process comprises the epiphytic growth stage and the subsequent penetration through the stomata, and requires warm temperatures and prolonged periods of high humidity or free moisture (Mondal and Timmer 2006; Timmer and Gottwald 2000). Warm humid nights are highly conducive for epiphytic growth and infection by *Z. citri-griseum* (Whiteside 1974). In Florida, infections can occur at any time and disease cycles overlap throughout the year (Timmer et al. 2004; Timmer and Gottwald 2000). In contrast, in Costa Rica, most of the infections occur at the beginning of the rainy season (Timmer and Gottwald 2000).

Considering that the knowledge on greasy spot epidemiology from other regions cannot be directly extrapolated to Panama, the objective of this study was to determine the dynamics of defoliation, inoculum production, airborne inoculum, and infection periods of *Z. citri-griseum* and their association with environmental conditions in Panama.

## 2 MATERIALS AND METHODS

### 2.1 Experimental orchard

The study was carried out in a commercial sweet orange ‘Valencia’ orchard severely affected by greasy spot in Churuquita Grande, Coclé province, Panama. The trees were 12 years old, grafted on ‘Swingle’ citrumelo (*C. paradisi* x *Poncirus trifoliata*) rootstock, and planted on a 3-by-7-m spacing. An experimental area of ≈0.17 ha where no fungicides were applied during the 2-year period of study from March 2011 to March 2013 was established in the center of the orchard. The presence of *Z. citri-griseum* in the orchard was confirmed by fungal isolation, morphological and molecular identification, and pathogenicity tests on sweet orange plants as previously described by Aguilera-Cogley et al. (2017).

Environmental data were monitored hourly in the orchard with an automated meteorological station (Watch Dog Series 1,000 Plant Disease Station, Spectrum Technologies, USA) including sensors for temperature (accuracies ± 0.6°C), relative humidity (accuracies ± 3.0%), rainfall (resolution 0.2 mm), and leaf wetness with a range of 0 (dry) – 15 (wet). Environmental monitors were located within the row in the experimental area at the site of a missing tree. Data were collected at 1.5 m above the soil surface, in the top quadrant of the canopy height. Leaf wetness sensors were placed with a northerly exposure and fixed at a 30-degree angle from the horizontal.

### 2.2 Defoliation dynamics

Defoliation dynamics of the trees in the experimental orchard were studied from March 2012 to April 2013. The trees were covered with a plastic net (10 × 10 m, 1-by-1-cm openings) fixed to the floor with ten stainless steel pins. Altogether, six groups of two trees were covered with a plastic net. Prior to the placement of the plastic net, all leaf litter present in each group of trees was eliminated. Then, newly defoliated leaves on the floor in each group of trees were collected monthly and weighed.

### 2.3 Inoculum dynamics

The dynamics of *Z. citri-griseum* inoculum production in the leaf litter was studied from March 2011 to March 2013. Leaf litter was collected monthly on the floor of the 24 trees selected and covered with a plastic mesh (2 × 2 m, 5-by-5-mm openings) fixed with four stainless steel pins. A sample of 25 g of leaf litter was collected from each plastic mesh every 15 days and 10 leaves were selected arbitrarily to estimate the degree of decomposition using a scale of 0 to 3, where 0 = not decomposed, still firm; 1 = partially decomposed, flexible still intact; 2 = moderately decomposed, some loss of lamina; and 3 = highly decomposed, skeletonized leaves (Mondal and Timmer 2002). The index of leaf decomposition (*ILD*) was calculated as:

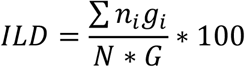

where *n*_*i*_ = number of units in category i, *g*_*i*_ = degree of category i, N = total number of units evaluated, and G = maximum degree of the scale. With the resulting data, the number of days until total leaf decomposition (*DLD*) was calculated when the sample reached the value of 100%. To quantify the number of ascospores released from leaf litter (*ASCL*), the total sample of 25 g of leaf litter was then soaked for 15 min in distilled water. Immediately after soaking, leaves were placed with the abaxial surface facing upward in a wind tunnel for 30 min until they were visibly dry (Vicent et al. 2011; Whiteside 1973). During the process, air and water temperature were maintained at ≈23°C. The ascospores released from leaf litter were collected on a glass microscope slide (26 × 76 mm) coated with silicone oil (Merck). Glass slides were stained with lactophenol-acid cotton blue and examined at a magnification of 400x. All ascospores showing the morphological characteristics of *Z. citri-griseum* described by Whiteside (1970) were counted in four microscope field transects. For each 15-day period, the following meteorological variables were measured: number of rainy days >1mm (*NRD*), accumulated rainfall (*AR*), average temperature (*AT*), and average relative humidity (*ARH*).

### 2.4 Airborne inoculum and infection periods

Airborne inoculum and infection periods of *Z. citri-griseum* were studied from March 2012 to April 2013. The dynamics of airborne ascospores (*AASC*) were studied by placing two glass microscope slides (26 × 76 mm) coated with silicone oil (Merck) in the center of the experimental area. The slides were placed under a plastic rain shelter (0.3 × 0.3 m) 0.25 m above the soil surface at a 45-degree angle from the horizontal (Campbell and Madden 1990). Microscope slides were changed weekly and treated with lactophenol-acid cotton blue, and a coverslip (32 × 22 mm) was affixed. Mountings were examined at a magnification of 400x and all ascospores with the morphological characteristics of *Z. citri-griseum* were counted as described above.

Infection periods were determined by placing a trap plant in the experimental area each week. Trap plants were 2-year sweet orange ‘Valencia’ trees grafted on ‘Swingle’ citrumelo grown in plastic pots (250 mm in diameter by 200 mm deep) containing potting mix (75% peat and 25% sand [vol/vol]). Trees were pruned completely to force growth of new shoots and were exposed weekly in the experimental orchard. Exposed trap plants were returned to a screenhouse, and disease incidence (*INC*) was evaluated after 10 months considering the total percentage of symptomatic leaves. For each weekly period, the following meteorological variables were measured: *NRD, AR, AT, ARH*, leaf wetness duration in hours (*LW*), number of periods of leaf wetness >1 hour per week (*PLW*), average duration of periods of leaf wetness >1 hour per week (*APLW*).

### 2.5 Data analyses

All data analyses were carried out with the software R version 4.1.0 (R Core Team 2022). Data on dynamics of defoliation, inoculum production, airborne inoculum, and infection periods were analyzed descriptively. Correlations among meteorological variables, inoculum production, airborne inoculum, and disease incidence on trap plants were analyzed with Pearson’s correlation coefficient. Pairwise correlations were classified as |r| ≤0.7 or |r| >0.7 according to Dormann et al. (2012), who proposed this threshold of correlation as an appropriate indicator for when collinearity begins to severely distort model estimation and subsequent prediction. General linear models (GLMs) were used to analyze the relationships of *DLD* or *ASCL* with meteorological variables *NRD, AR, AT*, and *ARH*. The model for the *DLD* was fitted with a Poisson regression, and log link function was used between the linear predictor and the response variable. In addition to the meteorological variables, *ASCL* was also included in the model as an explanatory variable:

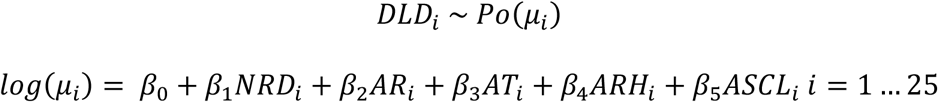

where *DLD* = number of days to total leaf decomposition, *β*_0_ and *β*_*j*_, *j* = 1,…, 5 are the parameters of the full model.

The model for *ASCL* was initially adjusted with a Poisson regression, but problems of overdispersion were found and the negative binomial distribution was used instead:

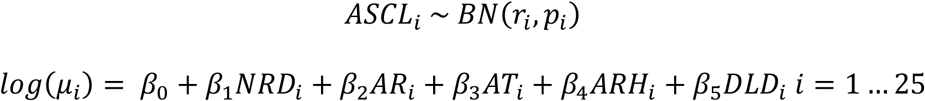

where *ASCL*_*i*_= number of ascospores released until its total release in *r*_*i*_ times in the month_i_. 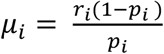 (*r*_*i*_ = the number of successes where all ascospores are released and *p*_*i*_= is the probability that ascospores are not released). *β*_0_ and *β*_*j*_, *j* = 1,…, 5 are the parameters of the full model.

Model selection was performed through a sequential search based on the lowest value of the Akaike Information Criterion (AIC) (Akaike 1974) following the principle of maximum parsimony. Additionally, the absence of overdispersion in the model was assessed with the dispersion parameter, dividing the residual deviance by the degrees of freedom. Diagnosis of the response variable *DLD* and *ASCL* was carried out by means of the graphic representation of the residual deviance.

GLMs were also used to analyze the relationships of *AASC* or *INC* with the meteorological variables. For each response variable, two types of models were proposed. The first model included only meteorological variables and the second also incorporated a temporal component. Due to the complexity of these models they were adjusted by means of Bayesian statistics with the Integrated Nested Laplace Approximation (INLA) (Rue et al. 2009). Unlike frequentist statistics, Bayesian statistics incorporates prior information through distributions of probability that, together with the likelihood function of the data, provide the a posteriori distribution of the parameters. In our study, the default non-informative prior distributions in the R-INLA package were applied. The models with the lowest value of the Deviance Information Criterion statistic (DIC) (Spiegelhalter et al. 2002) were selected. Additionally, the Conditional Predictive Ordinate (LCPO) (Roos and Held 2011), which evaluates the capacity to predictive the model was calculated as – 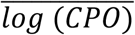. The best model was considered to be the one that had the LCPO value closest to zero.

The variable *AASC* was modeled using Poisson regression with the log link function between the linear predictor and the response variable. In the first model, *NRD, AR, AT*, and *ARH* were considered as explanatory variables. In the second model, an autoregressive temporal component of order 1 denoted *week* was added to the meteorological variables, so the number of ascospores could depend on the value of the previous week.

Model 1: Explanatory meteorological variables

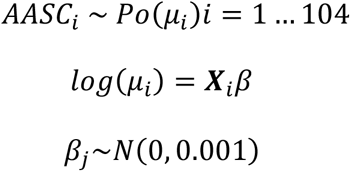

Model 2: Explanatory meteorological variables + temporal component, likelihood function:

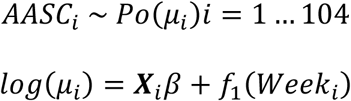

where:

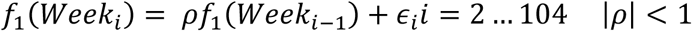

A priori distributions of the parameters:

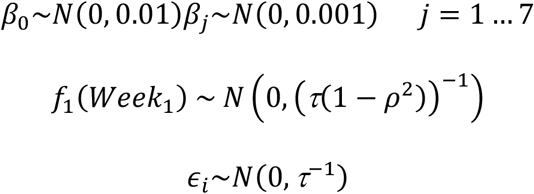

A priori distributions of the hyperparameters used by default R-INLA:

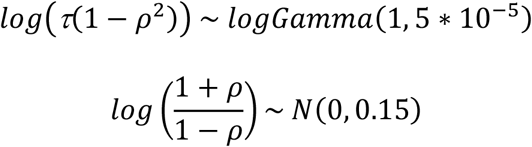

The variable *INC* was modeled by logistic regression using the logit link function between the linear predictor and the response variable. In the first model, *NRD, AR, AT, ARH, LW, PLW, APLW*, and *AASC* were considered as explanatory variables. In the second model, an autoregressive temporal component of order 1 denoted *week* was added, so the incidence of a week could depend on the value of the previous week.

Model 1: Explanatory meteorological variables

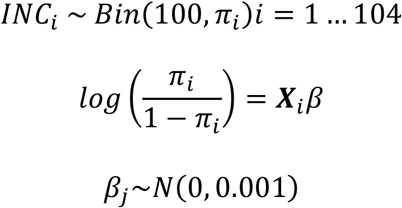

where:

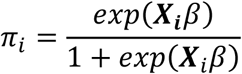

Model 2: Explanatory meteorological variables + temporal component, likelihood function:

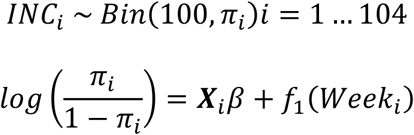

where:

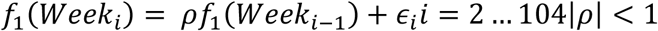

A priori distributions of the parameters:

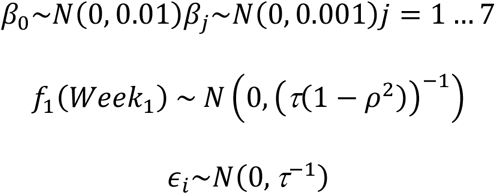

A priori distributions of the hyperparameters used by default R-INLA:

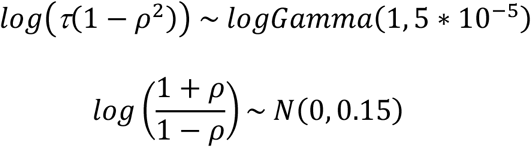

## 3 RESULTS

### 3.1 Defoliation dynamics

Two periods with different defoliation intensities were observed in the trees affected by greasy spot. One with low defoliation between May and November, with the lowest weight of leaf litter in June with 41.6 g tree^-1^ (Figure 1). The period from December to April was characterized by a greater defoliation, with January having the greatest amount of leaf litter with 697.5 g tree^-1^ (Figure 1).

**Figure 1.**
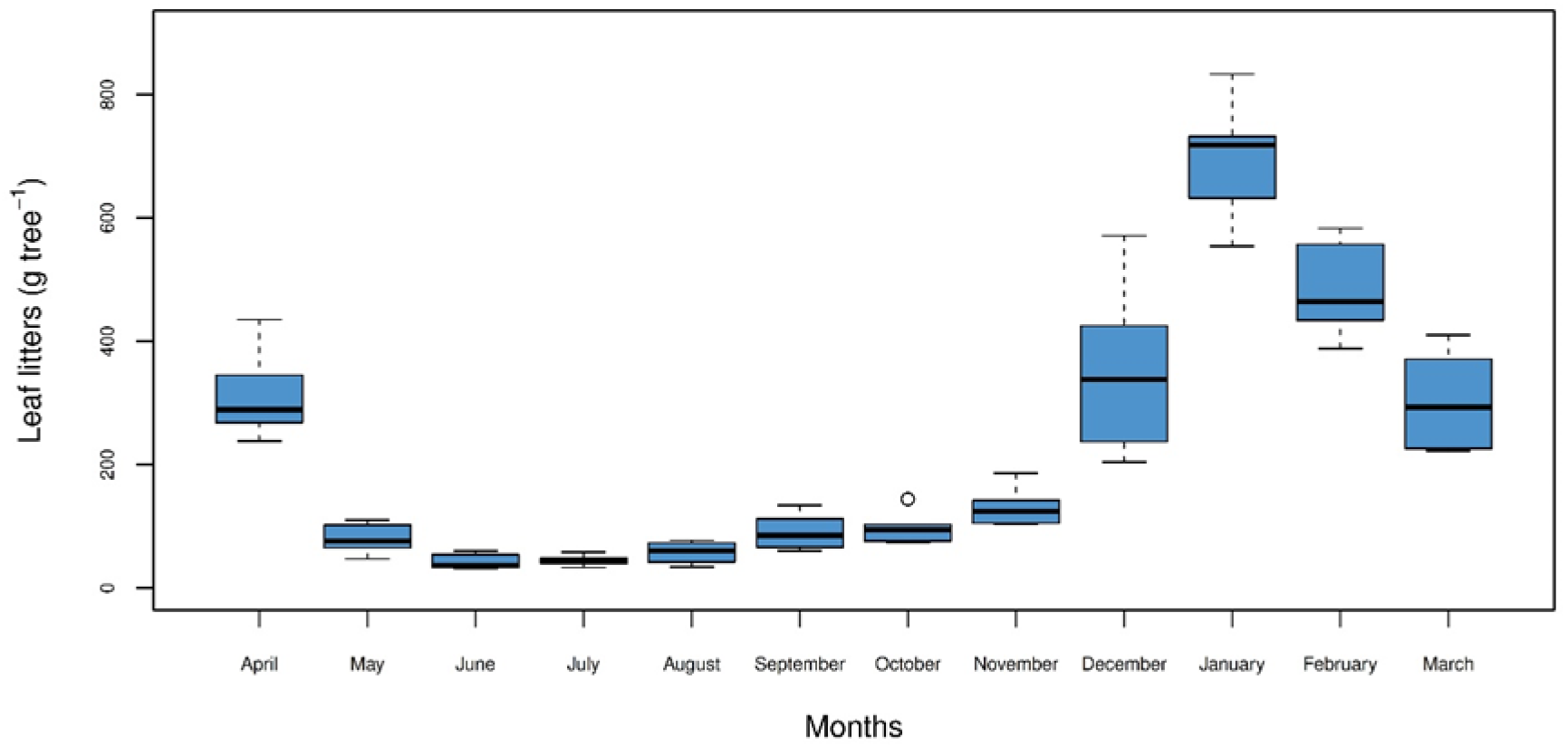
Defoliation dynamics in sweet orange ‘Valencia’ trees affected by citrus greasy spot from April 2012 to March 2013 in Churuquita Grande, Panama.

### 3.2 Inoculum dynamics

In the descriptive analysis, the variables *AT* and *ARH* presented lower variability than *ASCL* (Figure S1). Values of Pearson’s correlation coefficient for *DLD* varied from - 0.81 with *ARH* to 0.18 with *NRD* (Figure S2). For *ASCL*, the highest values were 0.54 and 0.53 with the *DLD* and *AT*, respectively, and the lowest was -0.01 with *AR* (Figure S2).

The model fitted using Poisson regression for *DLD* including the explanatory variables *NDR, AR*, and *ARH* (Table 1) showed the lowest AIC (163.33). The selection of the explanatory meteorological variables was threshold-based with pairwise correlations |r| ≤0.7. All these explanatory variables were significant (P<0.05) in the model. The assumption of the Poisson distribution of equality of mean and variance for the *DLD* variable was verified. The dispersion parameter was calculated by dividing the value of the residual deviance (24.40) of the model by 18 degrees of freedom, resulting in a value of 1.36, thus indicating the absence of over-dispersion. Model diagnosis was carried out by means of the graphic representation of the deviance of the residues, observing that they were concentrated within the values -2 and 2, with only one residue slightly above 2 (Figure S3A).

**Table 1.** Coefficients of the Poisson regression for the response variable number of days until total leaf decomposition (*DLD*).

| Parameters<sup>1</sup> | Estimate | se<sup>2</sup> | z-Value | *p*-Value |
| --- | --- | --- | --- | --- |
| Intercept | 12.603 | 0.739 | 17.057 | <0.0001 |
| *NRD* | 0.036 | 0.005 | 7.351 | <0.0001 |
| *AR* | -0.001 | 2.90e<sup>-03</sup> | -2.248 | 0.0246 |
| *ARH* | -0.111 | 0.009 | -11.697 | <0.0001 |

<sup>1</sup> Number of rainy days >1mm (*NRD*), accumulated rainfall (*AR*), average relative humidity (*ARH*).
<sup>2</sup> Standard error.

The resulting model indicated that for *NDR*, keeping the other variables constant, the average number of days until total leaf decomposition would increase by 3.67% for each day with >1mm rainfall (Table 1). For *AR*, keeping the other variables constant, the average number of days until total leaf decomposition would decrease by only 0.10% if accumulated rainfall increases by 1 mm (Table 1). With *ARH*, keeping the other variables constant, the average number of days until total leaf decomposition would decrease by 10.48% if the average relative humidity increased by 1% (Table 1).

The model fitted using the negative binomial regression for *ASCL* including the explanatory variables *NDR, AR, AT*, and *DLD* (Table 2) showed the lowest AIC (415.52). The selection of the explanatory meteorological variables was threshold-based with pairwise correlations |r| ≤0.7. All these explanatory variables were significant (P<0.01) in the model. The dispersion parameter calculated as the ratio between the residual deviance (29.95) and 20 degrees of freedom resulted in a value of 1.50, indicating the absence of over-dispersion. The deviance residues were concentrated between -2 and 2, with only one residue slightly above 2 (Figure S3B).

**Table 2.** Coefficients of the negative binomial regression for the response variable number of ascospores from leaf litter (*ASCL*).

| Parameters<sup>1</sup> | Estimate | se<sup>2</sup> | z-Value | p-Value |
| --- | --- | --- | --- | --- |
| Intercept | -1.030e<sup>+02</sup> | 1.614e<sup>+01</sup> | -6.383 | <0.0001 |
| *NRD* | -1.398e<sup>-01</sup> | 4.546e<sup>-02</sup> | -3.076 | <0.0001 |
| *AR* | 8.199e<sup>-03</sup> | 2.704e<sup>-03</sup> | 3.032 | <0.0001 |
| *AT* | 4.213e<sup>+00</sup> | 6.349e<sup>-01</sup> | 6.636 | <0.0001 |
| *DLD* | 5.324e<sup>-02</sup> | 9.893e<sup>-03</sup> | 5.382 | <0.0001 |

<sup>1</sup> Number of rainy days >1mm (*NRD*), accumulated rainfall (*AR*), average temperature (*AT*), number of days until total leaf decomposition (*DLD*).
<sup>2</sup> Standard error.

The resulting model indicated that for *NRD*, keeping the other variables constant, the number of ascospores released from leaf litter (*ASCL*) would decrease by 13.05% for each day with rainfall of >1mm (Table 2). With *AR*, keeping the other variables constant, the *ASCL* would increase by 0.82% for each mm increase in accumulated rainfall (Table 2). With *AT*, keeping the other variables constant, the relative risk of an increase in the *ASCL* is 4.21 times higher if the *AT* increases by 1°C (Table 2). Finally, for *DLD*, keeping the other variables constant, the *ASCL* would increase by 5.47% for each day until the total decomposition of the leaves (Table 2).

### 3.3 Airborne inoculum and infection periods

The variable *AASC* presented a seasonal behavior with a pattern of about 52 weeks (Figure 2). The value recorded at week 57 was somewhat out-of-range but it was decided to keep it in the analysis. Similarly, the variable *INC* presented a seasonal behavior with a pattern of about 52 weeks (Figure 2). The results observed clearly indicated that the value of the observation of *AASC* and *INC* in week *i* is likely related to the observation of the previous week *i*-1. This motivated the inclusion of a temporal component in these models to analyze the variables *AASC* and *INC* in a more adequate way.

**Figure 2.**
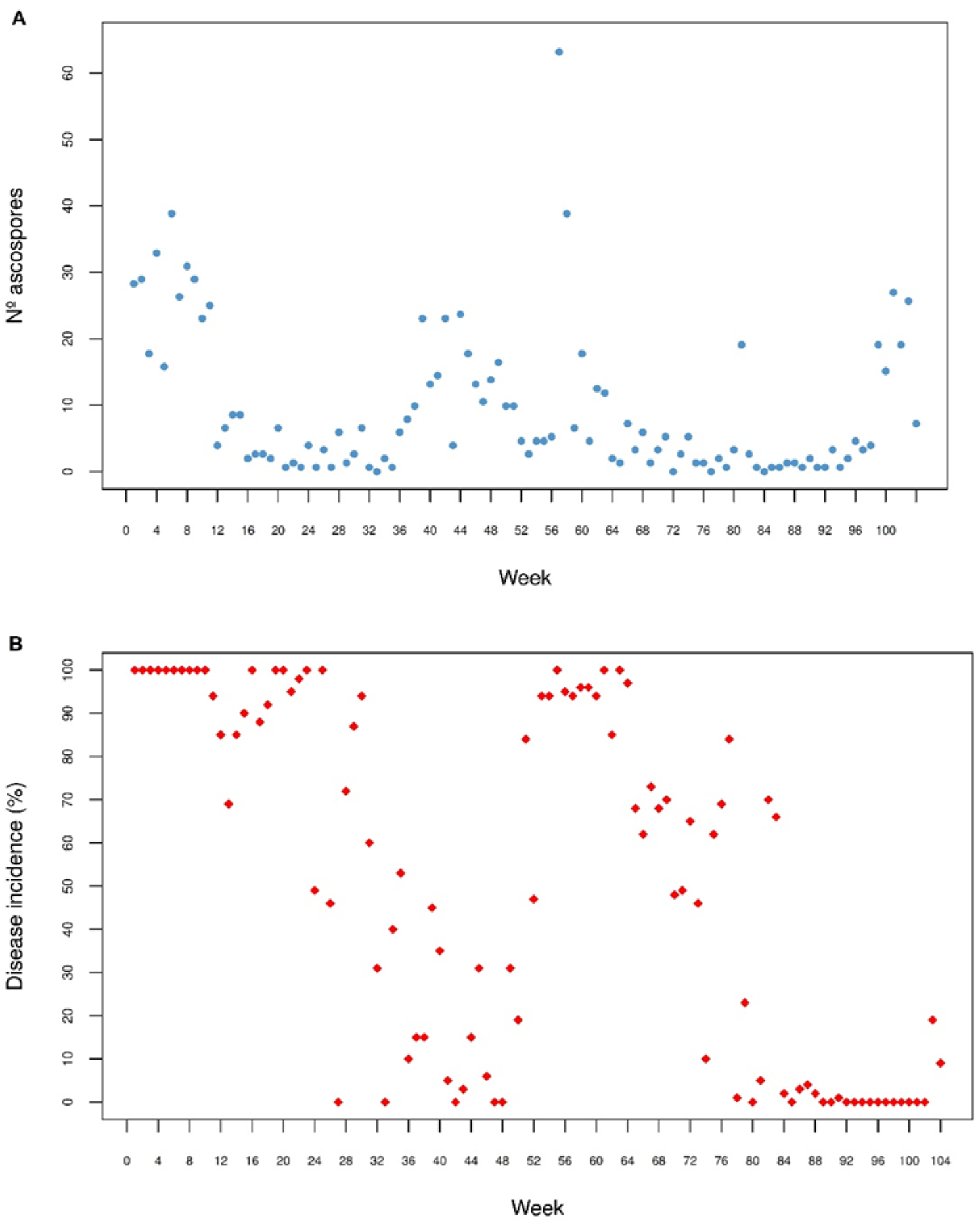
A) Weekly values of the number of airborne ascospores (*AASC*); B) disease incidence (*INC*) of *Zasmidium citri-griseum* from March 2011 to March 2013 in Churuquita Grande, Panama.

The box-and-whisker plots of the meteorological variables showed a very low variability of *AT* and higher for *AASC, INC, AR*, and *LW* (Figure S4A and S4B). The presence of ‘outliers’ was observed in the variable *AASC* and *AR*. The variables *NRD, ARH, PLW*, and *APLW* presented relatively symmetric distributions, with very close mean and median values (Figure S4A and S4B).

Pearson’s coefficients indicated a very low correlation between the meteorological variables and the variable *AASC*. The lowest value of the coefficient was 0.02 with *AT* and the highest was -0.44 with *ARH*. With the variable *INC*, the lowest value of the coefficient was -0.03 with *AT* and the highest was 0.33 with *APLW* (Figure S5). A high correlation was observed among the variables *LW, PLW*, and *APLW*, the highest being 0.92 between *LW* and *PLW* (Figure S5).

Table S1 shows the model of *AASC* with the lowest DIC, including only explanatory meteorological variables. In this case, the selected model did not include *AT*, which barely presented variability as can be observed in the box-and-whisker plots (Figure S4). Furthermore, *AT* and *AASC* presented the lowest linear correlation (Figure S5). Nevertheless, the predictive capacity of the model was not good, since the LCPO value was somewhat far away from 0 (Table S1).

By integrating an autoregressive temporal component of order 1 into the model of *AASC* through the variable *week*, it was possible to considerably reduce the DIC value and improve model fit. Table 3 shows the model for *AASC* with the lowest DIC of 524.66, including explanatory meteorological variables and the autoregressive temporal component of order 1. The selection of the explanatory meteorological variables was threshold-based with pairwise correlations |r| ≤0.7. The model selected included, in addition to the temporal component *week*, the explanatory meteorological variables *AR* and *NRD*. Nevertheless, the predictive capacity of the model was not good either, since LCPO (2.98) was much higher than 0.

**Table 3.** General linear model (GLM) for the number of airborne ascospores captured per week (*AASC*) with the explanatory meteorological variables and the autoregressive temporal component of order 1.

| Parameters and hyperparameters<sup>1</sup> | Mean | sd | $Q_{0.025}$ | $Q_{0.5}$ | $Q_{0.975}$ | Mode |
| --- | --- | --- | --- | --- | --- | --- |
| Intercept | 1.591 | 0.54 | 0.467 | 1.601 | 2.657 | 1.618 |
| *AR* | -0.006 | 0.002 | -0.01 | -0.006 | -0.002 | -0.006 |
| *NRD* | 0.106 | 0.056 | -0.005 | 0.106 | 0.216 | 0.105 |
| $\phi$ | 0.828 | 0.059 | 0.699 | 0.833 | 0.926 | 0.844 |
| $\tau$ | 0.756 | 0.231 | 0.372 | 0.734 | 1.268 | 0.692 |

<sup>1</sup> Mean, standard deviation (sd), quantiles ( $Q$ ), and mode for the parameters and hyperparameters. Accumulated rainfall (*AR*), and number of rainy days >1mm (*NRD*). $\phi$ is the dispersion parameter of the likelihood, and $\tau$ the precision of the random effect *week*. The 95% credible interval is defined by $Q_{0.025}$ and $Q_{0.975}$ .

The 95% credibility intervals for the relative risks of the model selected for *AASC* indicated that the number of airborne ascospores captured in a weekly period was related to the number in the previous week. Also, the relative risk of increasing the number of *AASC* would be greater if the number of rainy days >1mm (*NRD*) is greater but with less accumulated rainfall (*AR*) (Figure 3A).

**Figure 3.**
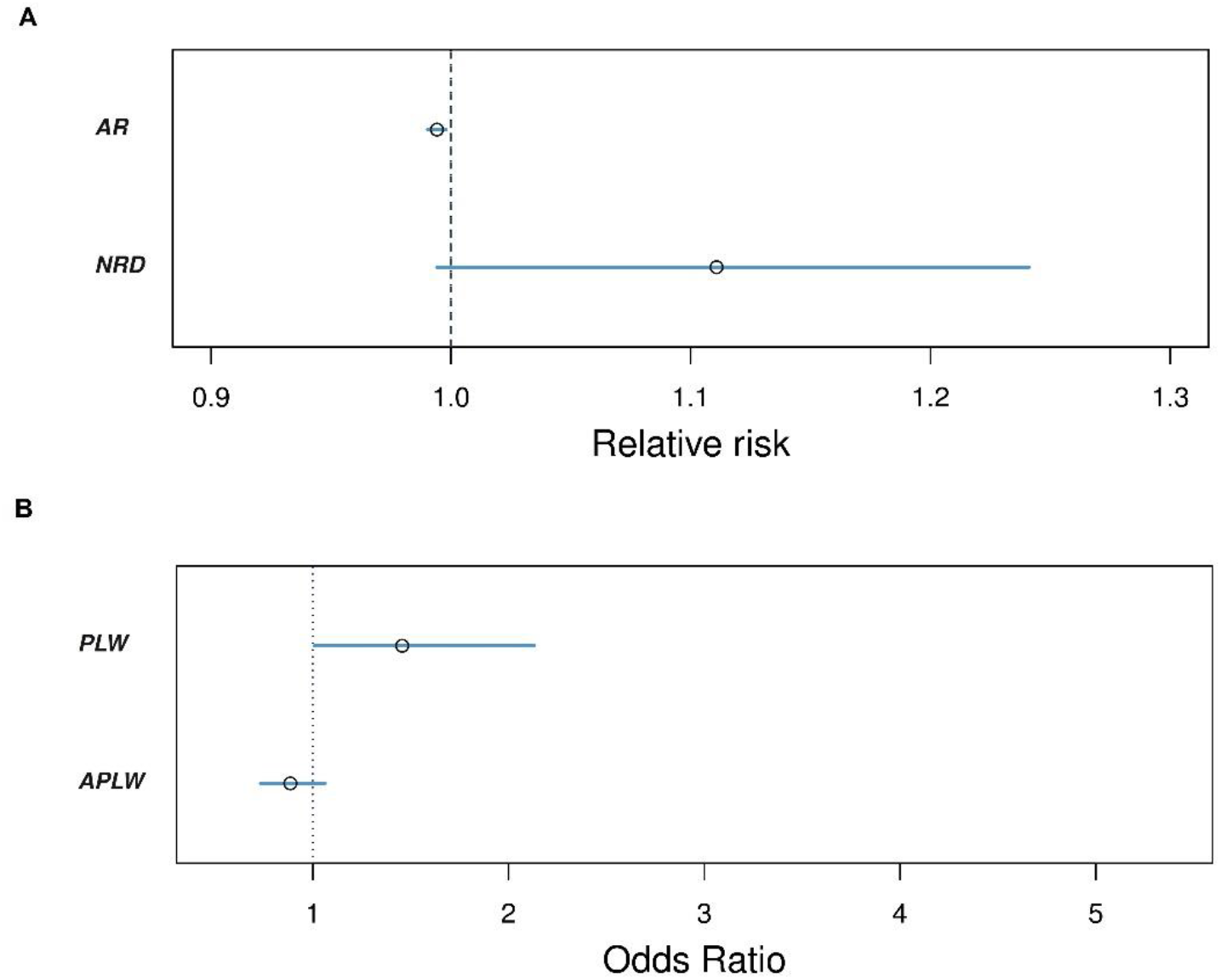
Graphic representation of the 95% credibility intervals: A) relative risk for the meteorological variables accumulated rainfall (*AR*) and number of rainy days >1mm (*NRD*) with the autoregressive temporal component of order 1 of the model for the number of airborne ascospores (*AASC*); B) odds ratio for the meteorological variables number of leaf wetness periods of >1 hour per week (*PLW*) and average duration of leaf wetness periods of >1 hour per week (*APLW*) with the autoregressive temporal component of order 1 of the model for disease incidence (*INC*).

Table S1 shows the model of *INC* with the lowest DIC, including only explanatory meteorological variables. The selected model did not include the meteorological variable *AT*, which presented little variability (Figure S4). Furthermore, *AT* and *INC* presented the lowest linear correlation (Figure S5). Nevertheless, the predictive capacity of the model was not good, since the LCPO value was rather far away from 0 (Table S1).

By integrating an autoregressive temporal component of order 1 into the model of *INC* through the variable *week*, it was possible to considerably reduce the value of DIC and improve model fit. Table 4 shows the model of *INC* with the lowest DIC (462) including the explanatory meteorological variables and the autoregressive temporal component of order 1. The selection of the explanatory meteorological variables was threshold-based with pairwise correlations |r| ≤0.7. The model selected included, in addition to the temporal component *week*, the explanatory meteorological variables *PLW* and *APLW*. Nevertheless, the predictive capacity of the model was not good either, since LCPO (15.94) was much higher than 0.

**Table 4.** General linear model (GLM) for disease incidence (*INC*) with the explanatory meteorological variables and the autoregressive temporal component of order 1.

| Parameters and hyperparameters<sup>1</sup> | Mean | sd | $Q_{0.025}$ | $Q_{0.5}$ | $Q_{0.975}$ | Mode |
| --- | --- | --- | --- | --- | --- | --- |
| Intercept | -0.585 | 3.35 | -7.549 | -0.627 | 6.664 | -0.673 |
| *PLW* | 0.39 | 0.193 | 0.017 | 0.387 | 0.775 | 0.383 |
| *APLW* | -0.117 | 0.098 | -0.312 | -0.116 | 0.073 | -0.114 |
| $\phi$ | 0.854 | 0.04 | 0.762 | 0.859 | 0.919 | 0.868 |
| $\tau$ | 0.038 | 0.013 | 0.018 | 0.037 | 0.068 | 0.033 |

<sup>1</sup> Mean, standard deviation (sd), quantiles ( $Q$ ), and mode for the parameters and hyperparameters. Number of leaf wetness periods of >1 hour per week (*PLW*); and average duration of leaf wetness periods of >1 hour per week (*APLW*). $\phi$ is the dispersion parameter of the likelihood and $\tau$ is the precision of the random effect *week*. The 95% credible interval is defined by $Q_{0.025}$ and $Q_{0.975}$ .

The 95% credibility intervals for the odds ratio of the model selected for *INC* indicated that the disease incidence in a weekly period depends on that of the previous period. Also, the presence of shorter but more frequent periods of leaf wetness >1 hour (*PLW*) would increase disease incidence (Figure 3B).

## 4 DISCUSSION

In the study of the dynamics of defoliation of ‘Valencia’ sweet orange trees affected by greasy spot, the greatest leaf litter amounts were generated from December to April (Figure 1). In Panama, these months are characterized by being relatively dry with low rainfall of around 100 mm per month (WBG 2022). Citrus orchards in Panama are seldom irrigated and so the water stress probably favors greater defoliation during this period in the trees affected by greasy spot. In Florida, winter and spring are relatively dry and leaf litter is mainly produced from late January to April (Timmer and Gottwald 2000). Similar results were found in our study, where the period with the greatest defoliation of the trees coincided with the dry months from December to April. However, not all defoliation can be attributed to the effect of greasy spot since citrus trees also have a natural loss of leaves. Following previous works (Aguilera-Cogley and Vicent 2020), it would be advisable to carry out experiments comparing trees treated and not treated with fungicides to determine the effect of greasy spot on defoliation more precisely.

The model for the response variable number of days until total leaf decomposition (*DLD*), included the explanatory variables number of rainy days >1mm (*NRD*), accumulated rainfall (*AR*), and average relative humidity (*ARH*) (Table 1). The coefficient for the variable *NRD* indicated that keeping the other variables constant, *DLD* would increase by 3.67% for each day with rainfall >1mm (Table 1). Experiments conducted in Florida demonstrated that relatively long wet periods of 3 to 5 days per week accelerate leaf litter decomposition compared with 1 to 2 days per week (Mondal and Timmer 2002). In fact, in most plant species, the rate of leaf litter decomposition increases with temperature and rainfall (Zhang et al. 2008). The results obtained in our experiment seem to contradict these studies. However, it must be taken into account that the variable *NRD* considers any rainfall >1mm, which also includes days with very low amounts of rainfall and may not have any influence on the process of leaf litter decomposition. In Panama, these low-volume rains between 1-10 mm day^-1^ are frequent during the dry season. The longer the leaves remain on the orchard floor (i.e., greater *DLD*), the more the accumulated number of days with this type of low intensity rainfall would also increase, hence the positive relationship between the two variables.

For the variable *AR*, the model indicated that keeping the other variables constant, the variable *DLD* would decrease slightly if accumulated rainfall increases. This is in line with field experiments carried out in Florida, where the increase in accumulated rainfall favored the rapid decomposition of citrus leaves in summer (Mondal and Timmer 2002). In Panama, the rainy season occurs from May to November with an average monthly rainfall over 220 mm, likely favoring the rapid decomposition of the leaves (WBG 2022). In contrast to *NRD*, the variable *AR* considers the total accumulated precipitation and it is a more adequate indicator of the amount of rainfall to which the leaf litter was exposed on the orchard floor.

The variable *ARH* showed the greatest variability in the model, indicating that *DLD* would decrease by 10.48% if *ARH* increases by 1% (Table 1). Experiments under controlled conditions with grapefruit leaves evaluated high relative humidity close to 100% with temperatures between 28°C and 32°C, resulting in a faster leaf litter decomposition than occurs at 20°C and 24°C (Mondal and Timmer 2002). In our study, the mean *ARH* was 82%, while the *AT* registered was 25°C (Figure S1D). These values are similar to those reported by Mondal and Timmer (2002), which indicates that leaf litter decomposition is favored by high relative humidity.

In the present study, the identification of the ascospores of *Z. citri-griseum* was based on their morphological characteristics. Nevertheless, this method might not be sufficiently specific considering other Mycosphaerellaceae coexisting in the leaf litter. The presence of *Z. citri-griseum* in the experimental orchard was confirmed by isolation and molecular methods (Aguilera-Cogley et al. 2017), but it would be necessary to develop, validate, and implement molecular methods for the specific detection and quantification of *Z. citri-griseum* in aerobiological monitoring systems. Currently, this aspect remains unresolved.

The model for the response variable number of ascospores released from leaf litter (*ASCL*) included the explanatory variables *DLD, NRD, AR*, and *AT* (Table 2). For the variable *DLD*, the model indicated that, keeping the other variables constant, *ASCL* would increase if the period of *DLD* is prolonged (Table 2). This could be attributed to the fact that a longer duration of the leaves on the orchard floor implies a greater opportunity for the formation and maturation of pseudothecia and therefore an increase in the number of ascospores produced.

For the variable *NRD*, the model indicated that, keeping the other variables constant, *ASCL* will decrease 13.05% if the number of days with rainfall >1mm increases (Table 2). In experiments conducted under controlled conditions, the highest number of ascospores released was achieved by wetting the leaves once (Mondal et al. 2003). However, when the leaves were moistened successively for several days, the amount of ascospores released decreased considerably compared to the initial day.

For the variable *AR*, the model indicated that, keeping the other variables constant, *ASCL* increased by only 0.82% for each millimeter of accumulated rainfall (Table 2). Therefore, the magnitude of this effect on the release of ascospores would be directly proportional to the rainfall accumulated during the period of permanence of the leaves on the orchard floor. In relation to the variable *AT*, which contributed with the greatest variability to the model with a relative risk of 4.21, indicated that *ASCL* increased with the average temperature during the period of permanence of the leaves on the orchard floor (Table 2). It has been reported that the greatest amounts of ascospores of *Z. citri-griseum* released from the leaf litter occur with a temperature range of 28-30°C (Mondal and Timmer 2002). Similar temperatures were recorded in our study (Figure S1D), with an average of 25°C that did not vary substantially throughout the year, probably favoring a constant release of ascospores.

In the descriptive analysis of the airborne inoculum of *Z. citri-griseum* (Figure 2A), the number of airborne ascospores captured showed the highest peaks in April and May, coinciding with the onset of the rainy season and the citrus leaf flush. In Costa Rica, the peak of *Z. citri-griseum* ascospores captured was in June, also coinciding with the beginning of the rainy season (Hidalgo et al. 1997). In Florida, the highest number of available ascospores also occurs in April and May (Timmer and Gottwald 2000; Timmer et al. 1995). In our study, the method of capturing ascospores on glass slides made it possible to determine the moments of the greatest amount of airborne inoculum. As previously mentioned, ascospores from other Mycosphaerellaceae species might be captured inadvertently. However, the study was conducted in an orchard severely affected by greasy spot, where the presence of *Z. citri-griseum* was confirmed by isolation and molecular methods. Moreover, the experimental area was large enough to avoid interferences from exogenous fungal inoculum coming from neighboring plant sources.

The model selected for the variable number of airborne ascospores (*AASC*) incorporates the effect of the autoregressive temporal component of order 1, together with the explanatory meteorological variables *NRD* and *AR* (Table 3). The autoregressive temporal component indicated that the number of airborne ascospores present in a week was related to the number of the previous week. This temporal autocorrelation indicates that the behavior of *Z. citri-griseum* ascospores in the air does not undergo abrupt changes from one week to the next, but rather evolves smoothly in an increasing or decreasing way depending on the time of year.

As regards the effect of the *NRD*, the model indicated that the number of airborne ascospores would increase with the number of days with rainfall >1mm. In the two years analyzed in the study, the peaks of airborne ascospores were preceded by relatively dry periods, when the leaf litter decomposes more slowly. The frequency of rainfall during the dry season is typically low, favoring a situation in which the leaf litter is wet in some moments but then dries quickly. These conditions may favor the formation and maturation of pseudothecia in the leaf litter, which will release the ascospores at the onset of the rainy season. In Florida, alternating wet and dry periods accelerate the formation of the pseudothecia of *Z. citri-griseum* on the decomposing leaves (Whiteside 1970).

In relation to the variable *AR*, the model indicated that the number of airborne ascospores captured would decrease with the volume of accumulated rain. In Panama, rain is most frequent from May to November, with monthly rainfall averages of about 290 mm (WBG 2022). These climatic conditions would favor a rapid decomposition of the leaf litter, limiting the formation and maturation of pseudothecia and thus reducing the production of ascospores (Mondal and Timmer 2002).

With the disease incidence (*INC*) of greasy spot in the trap plants, variations were observed during the two years analyzed in the study. The highest *INC* values were observed at the beginning of the rainy season, from the end of April to June, coinciding with the maximum values of airborne ascospores. Infections were also observed throughout the year, albeit with lower *INC* values. In Florida, if environmental conditions are favorable and inoculum is available, infections by *Z. citri-griseum* can occur at practically any time of the year (Timmer et al. 1995). Experiments also conducted in Florida indicated that greasy spot severity on trap plants can be moderate to high throughout the year, except in the cooler months from November to February (Timmer et al. 2000).

The model selected for *INC* integrated additively the effect of the autoregressive temporal component of order 1 and the explanatory meteorological variables number of periods of leaf wetness >1 hour per week (*PLW*) and average duration of leaf wetness periods of >1 hour per week (*APLW*) (Table 4). As in the case of the number of airborne ascospores (*AASC*), the autoregressive temporal component indicated that the incidence of greasy spot in the trap plants in one week depends on that of the previous week. The model also incorporated the variables *PLW* and *APLW*, indicating that short and frequent wetting periods would favor infections by *Z. citri-griseum* in trap plants. In Florida, leaf wetness periods of 6 h or longer during the night favored the development of the epiphytic mycelium and infection by *Z. citri-griseum* (Whiteside 1974). In our study, the duration of leaf wetness periods was 11 h on average (Figure S4B), which, together with the high relative humidity (Figure S4B), was conducive to the development of epiphytic mycelium on the leaves and subsequent infections by *Z. citri-griseum*.

The resulting models for the variables number of airborne ascospores captured (*AASC*) and disease incidence (*INC*) presented low predictive capacity. This could be attributed to the relatively short duration of the study period and the resulting limited amount of data, as time series analyses require more years to obtain more robust models. In any case, it was possible to identify some climatic factors related to the presence of airborne ascospores and infections in trap plants. More significantly, our models highlighted the importance of temporal autocorrelation, an aspect often overlooked when analyzing data from trap plants and spore monitoring devices. The epidemiological information obtained in our study will allow more efficient strategies to be designed to control citrus greasy spot in Panama.

## ACKNOWLEDGEMENTS

VAC was supported by SENACYT through the National Research System (SNI) (grant 39-2021).

## DATA AVAILABILITY STATEMENT

The data that support the findings of this study are available from the corresponding author upon reasonable request.

## SUPPORTING INFORMATION

**Table S1.** General linear model (GLM) for the number of airborne ascospores captured per week (*AASC*) and disease incidence (*INC*) with the explanatory meteorological variables.

| Models<sup>a</sup> | DIC<sup>b</sup> | LCPO<sup>c</sup> |
| --- | --- | --- |
| *AASC* |  |  |
| Intercept + *AR* + *NRD* + *ARH* + *LW* + *PLW* + *APLW* | 1,350.14 | 6.85 |
| *INC* |  |  |
| Intercept + *AASC* + *AR* + *NRD* + *ARH* + *LW* + *PLW* + *APLW* | 7,415.81 | 38.09 |

<sup>a</sup> Airborne ascospores (*AASC*); disease incidence (*INC*); accumulated rainfall (*AR*); number of rainy days >1mm (*NRD*); average relative humidity (*ARH*); leaf wetness duration in hours (*LW*); number of leaf wetness periods of >1 hour per week (*PLW*); and average duration of leaf wetness periods of >1 hour per week (*APLW*).
<sup>b</sup> DIC: Deviance information criterion.
<sup>c</sup> LCPO: Logarithmic conditional predictive ordinate.

**Figure S1.**
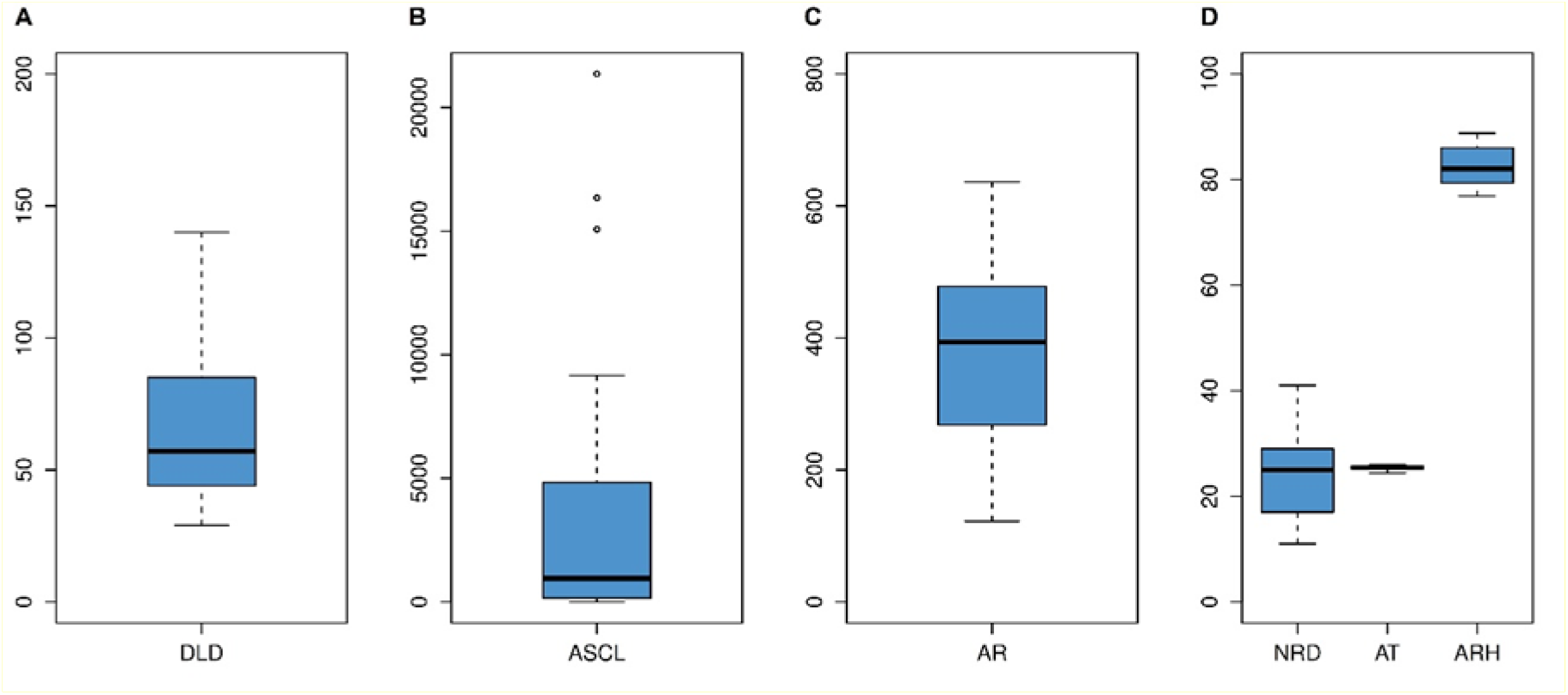
Box-and-whisker plots of the variables: A) number of days until total leaf decomposition (*DLD*); B) number of ascospores from leaf litter (*ASCL*); C) accumulated rainfall (*AR*); D) number of rainy days >1mm (*NRD*); average temperature (*AT*); and average relative humidity (*ARH*).

**Figure S2.**
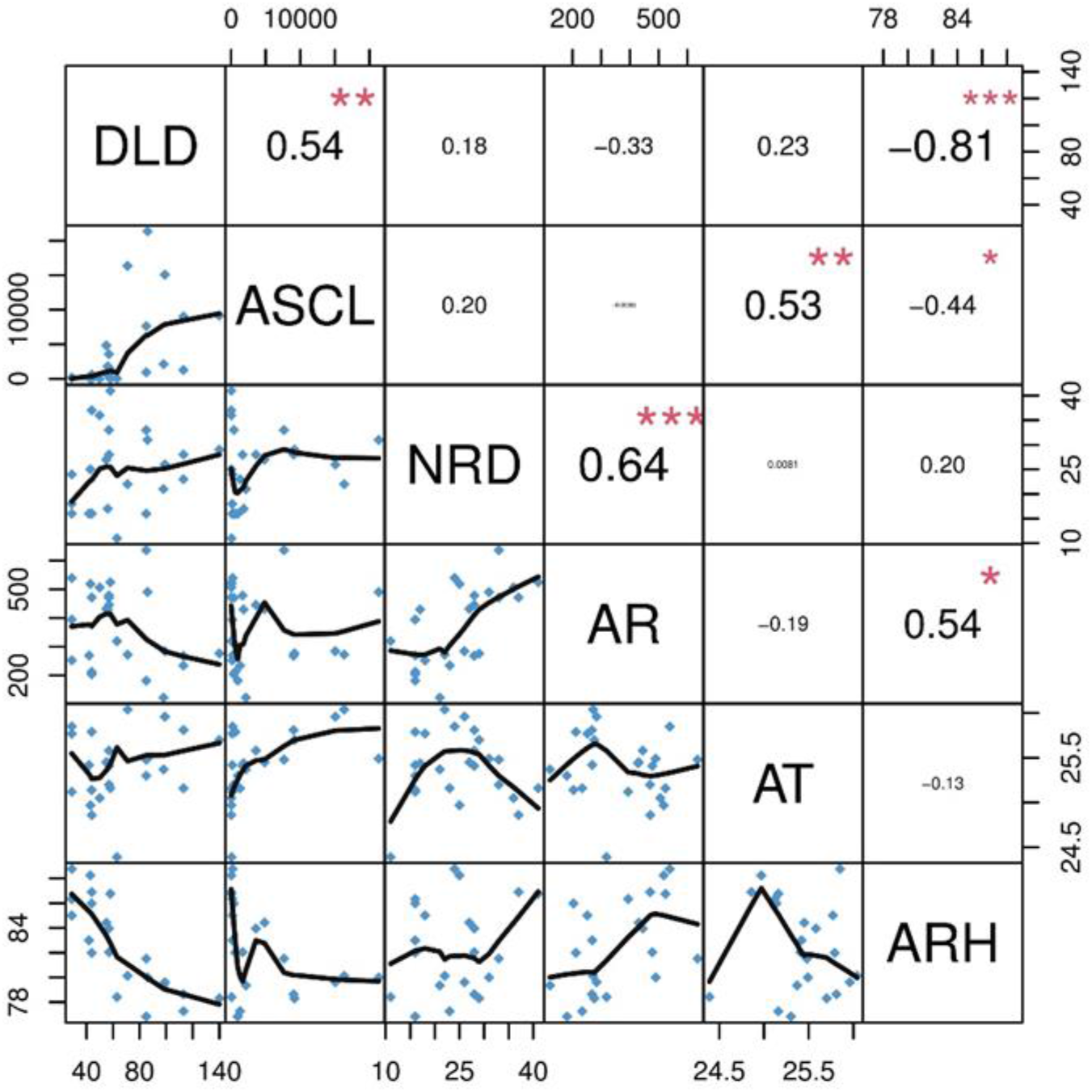
Correlation matrix for the variables number of days until total leaf decomposition (*DLD*); number of ascospores from leaf litter (*ASCL*); number of rainy days >1mm (*NRD*); accumulated rainfall (*AR*); average temperature (*AT*); and average relative humidity (*ARH*).

**Figure S3.**
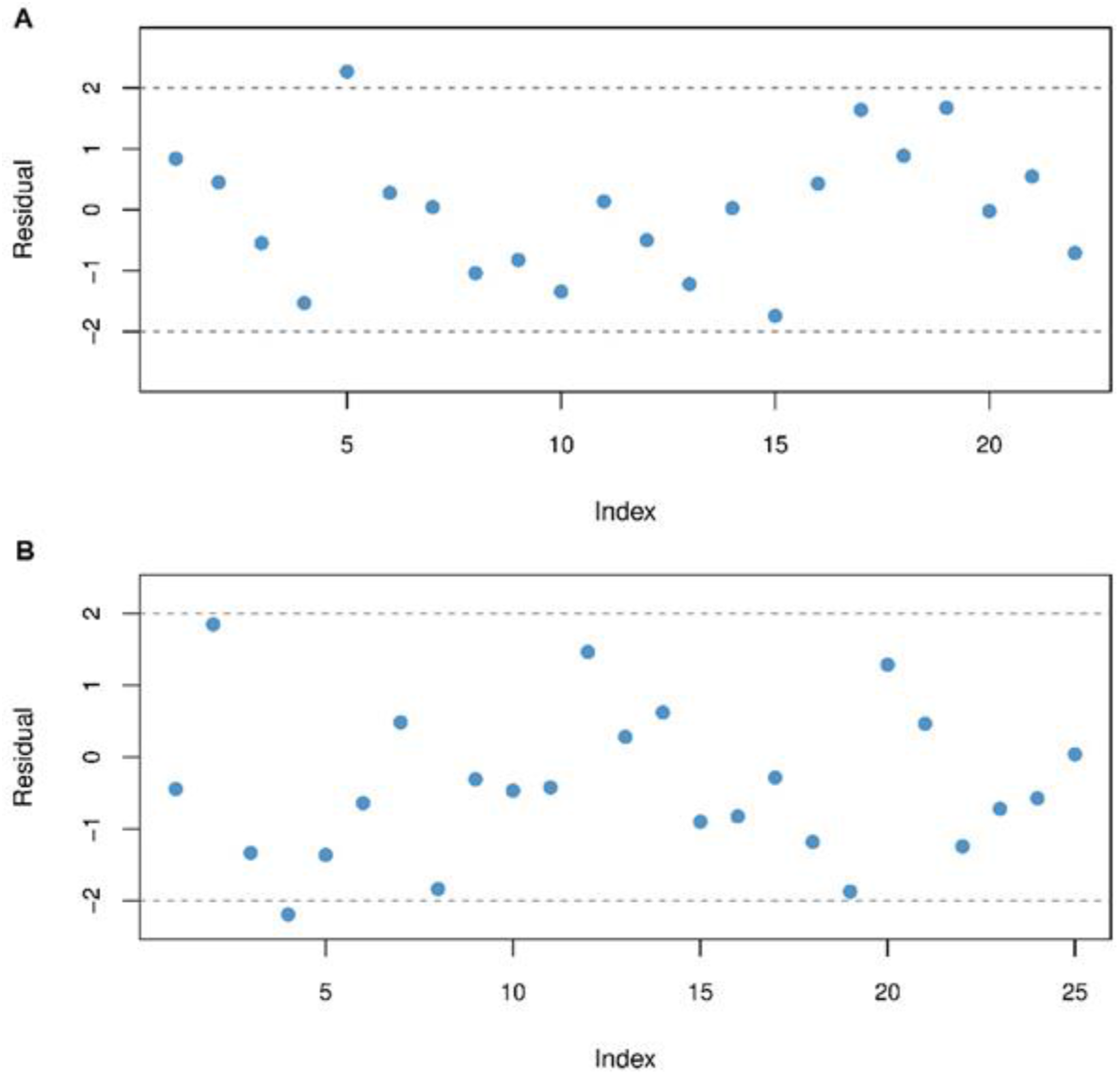
Graph of the deviance residuals of the general linear model (GLM) for: A) number of days until total leaf decomposition (*DLD*); B) number of ascospores from leaf litter (*ASCL*).

**Figure S4.**
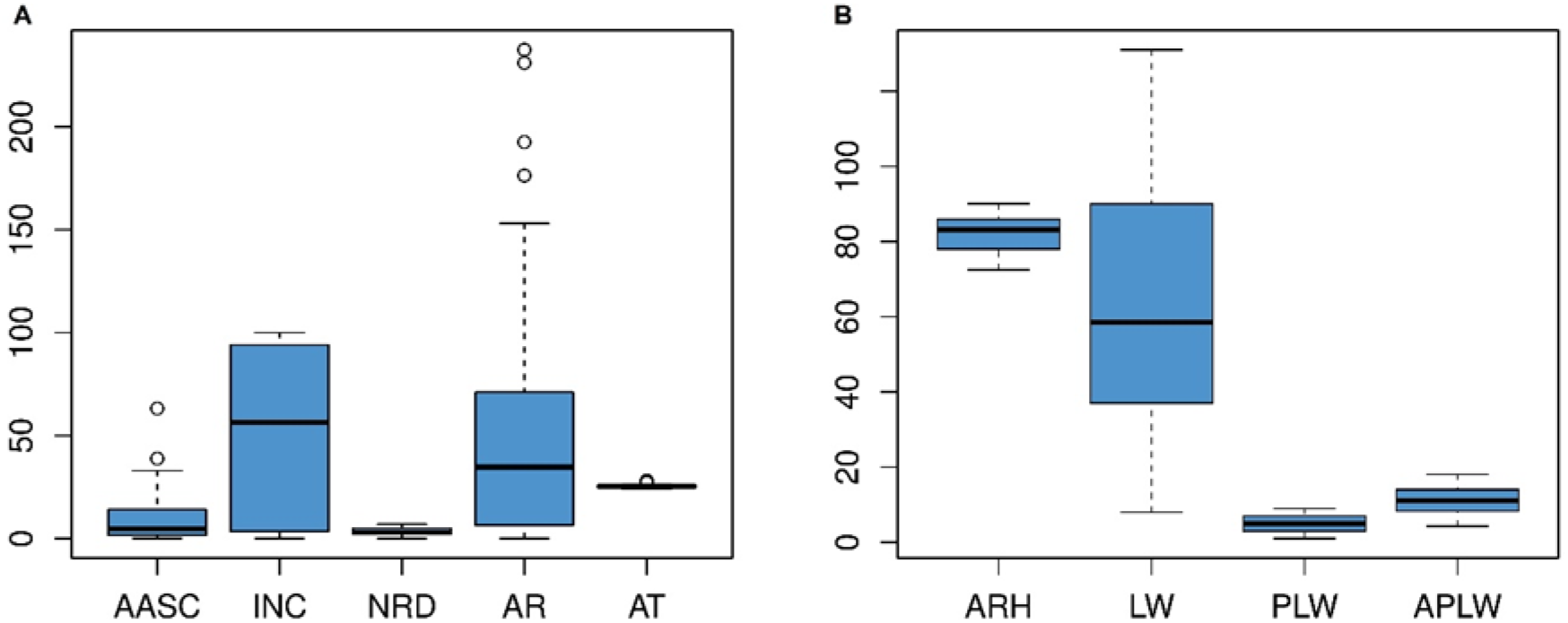
Box-and-whisker plots of: A) number of airborne ascospores (*AASC*); disease incidence (*INC*); number of rainy days >1mm (*NRD*); and accumulated rainfall (*AR*); average temperature (*AT*); B) average relative humidity (*ARH*); leaf wetness duration in hours (*LW*); number of leaf wetness periods of >1 hour per week (*PLW*); and average duration of leaf wetness periods of >1 hour per week (*APLW*).

**Figure S5.**
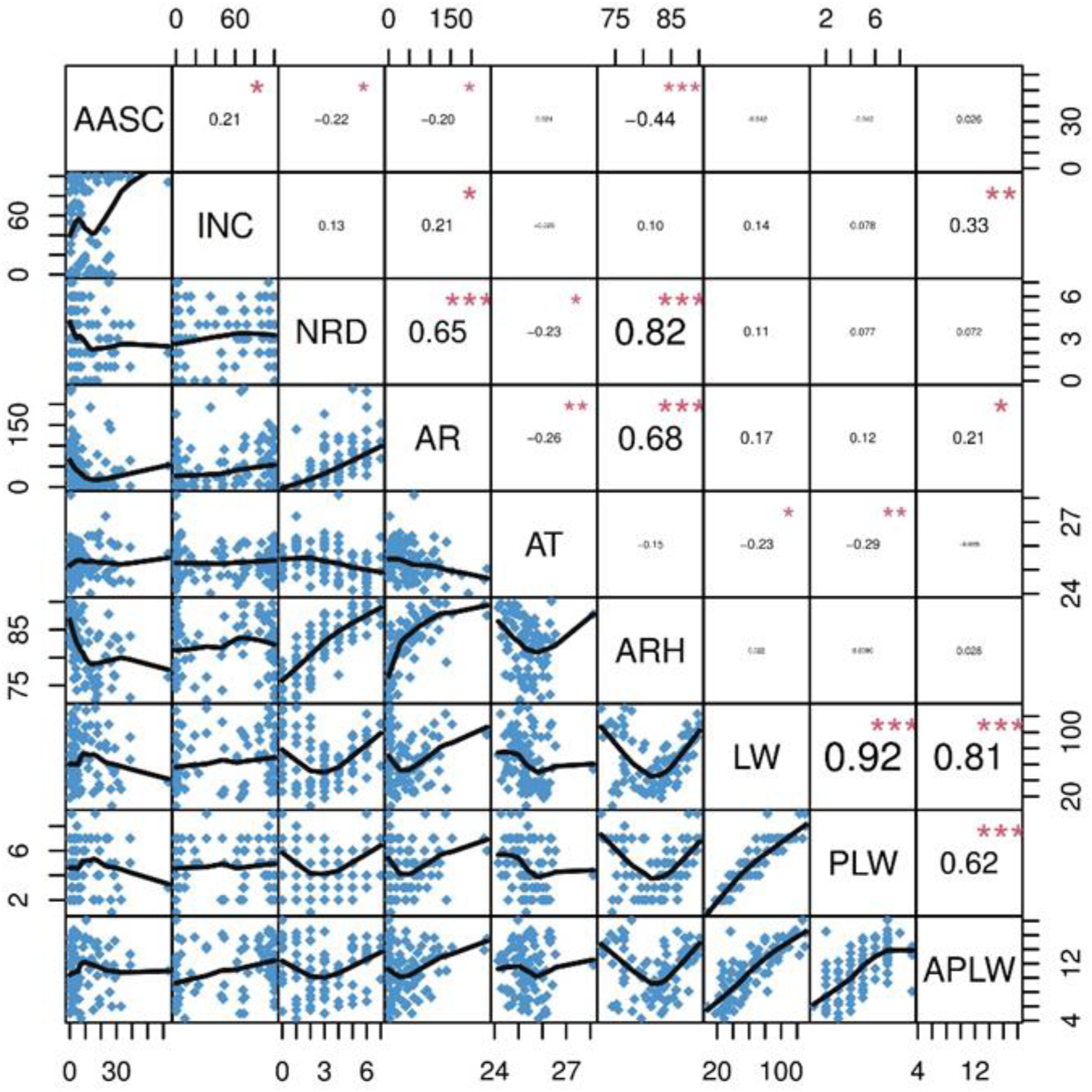
Correlation matrix for the variables: airborne ascospores (*AASC*); disease incidence (*INC*); number of rainy days >1mm (*NRD*); accumulated rainfall (*AR*); average temperature (*AT*); average relative humidity (*ARH*); leaf wetness duration in hours (*LW*); number of leaf wetness periods of >1 hour per week (*PLW*); and average duration of leaf wetness periods of >1 hour per week (*APLW*).

